# Cytoplasmic DNA Sensing Links LINE-1 Expression to Neuronal Senescence in Alzheimer’s Disease

**DOI:** 10.64898/2026.07.27.740588

**Authors:** Joseph R. Herdy, Emma E. Taylor, Lukas Karbacher, Oliver Borgogno, Larissa Traxler, Jessica Lagerwall, Vincent A. Huynh, Zornitsa V. Lefterova, Austin Kang, Carlota Tosat-Bitrian, Maxfield Kelsey, John Sedivy, Subhash Sinha, Li Gan, C. Frank Bennett, Dylan A. Reid, Jerome Mertens, Fred H. Gage

## Abstract

Cellular senescence contributes to neurodegeneration in Alzheimer’s disease (AD), yet brain-penetrant senotherapeutic strategies remain limited. Here, we identify long interspersed nuclear element 1 (LINE-1) retrotransposons as key regulators of neuronal senescence and the senescence-associated-secretory-phenotype (SASP) in AD. Using transdifferentiated induced neurons (iNs) that preserve donor-specific aging-associated molecular signatures, we show that pharmacological inhibition of LINE-1 with nucleoside reverse transcriptase inhibitors (nRTIs) or antisense oligonucleotides reduces p16 expression, suppresses SASP and interferon-stimulated gene programs, and attenuates paracrine induction of reactive astrogliosis. Spatial transcriptomic analysis of human AD brain tissue further supports that senescent neurons with high LINE-1 expression are localized to inflammatory niches in the brain. Although bulk analysis finds no significant differences in LINE-1 expression between AD and control neurons, long-read single-cell RNA sequencing of iNs identifies a subset of neurons with elevated LINE-1 activity which display transcriptional signatures of neurodegeneration, immune activation, and senescence are enriched in AD relative to controls. RNA velocity analysis indicates that LINE-1 activation precedes the induction of canonical senescence markers, supporting a causal rather than consequential role. Mechanistically, LINE-1-derived cytoplasmic DNA activates the cGAS-STING innate immune pathway in post-mitotic neurons, and inhibition of cGAS phenocopies the effects of LINE-1 suppression. Together, these findings establish a LINE-1/cGAS-STING axis as a driver of neuronal senescence in AD and highlight LINE-1 inhibition as a tractable senomorphic strategy for neurodegenerative disease.

## Introduction

Cellular senescence is increasingly recognized as a fundamental driver of aging and age-associated pathologies, including neurodegenerative disease^123^. Initially described as an irreversible cell-cycle arrest in proliferative cells, senescence is now understood as a complex stress response characterized by chromatin remodeling, metabolic dysfunction, and the acquisition of a pro-inflammatory senescence-associated secretory phenotype (SASP) that can disrupt tissue homeostasis ^4–6^. While senescence has historically been studied in dividing cell populations, recent work has established that post-mitotic cells, including neurons, can adopt senescence-like states. These “post-mitotic senescence” programs share key features with canonical senescence, including upregulation of cyclin-dependent kinase inhibitors such as p16 and p19, activation of DNA damage responses, and secretion of pro-inflammatory mediators ^7–11^. In neurons, such states have been linked to aging-associated dysfunction, tau pathology, and neurodegeneration ^9,12–15^. Importantly, the neuronal SASP in the brain can propagate dysfunction through paracrine signaling, inducing reactive states in neighboring glia and amplifying neuroinflammation ^16,17^.

A major barrier to studying neuronal aging has been the loss of age-associated features during reprogramming through induced pluripotent stem cell (iPSC) intermediates. Direct lineage conversion strategies circumvent this limitation by preserving donor age signatures. Notably, we and others have demonstrated that transdifferentiated induced neurons (iNs) retain key hallmarks of aging, including epigenetic, transcriptomic, and functional features that are erased during iPSC reprogramming ^18–25^. These models natively exhibit neuronal senescence in vitro without the need for exogenous or genetic stressors, facilitating investigation of upstream drivers and therapeutic interventions. Despite this, the initiating mechanisms of neuronal senescence remain incompletely understood.

One emerging mechanism linking aging to senescence and inflammation is the reactivation of transposable elements, particularly long interspersed nuclear element-1 (LINE-1). LINE-1 elements comprise approximately 17–20% of the human genome, although only a small subset of full-length elements remain retrotransposition-competent ^26,27^. LINE-1 expression is normally tightly suppressed through epigenetic mechanisms including DNA methylation and heterochromatin formation; however, this repression erodes with age and cellular stress ^28–30^. In multiple cell types, activation of LINE-1 has been shown to drive inflammatory responses through the generation of cytoplasmic nucleic acids that engage innate immune pathways ^28,29,31,32^. In the brain, LINE-1 activity is particularly notable due to its dynamic regulation in neurons and high levels of activity relative to other somatic tissues ^33–35^. A key mechanistic link between LINE-1 activity and inflammation is the generation of cytoplasmic DNA intermediates during reverse transcription. These nucleic acids can activate the cGAS-STING pathway, a central mediator of innate immune responses to aberrant DNA species ^36,37^. Activation of cyclic GMP-AMP synthase (cGAS) by cytoplasmic DNA leads to stimulation of STING and downstream induction of type I interferon signaling. In senescent cells, cytoplasmic chromatin fragments and retrotransposon-derived DNA have been shown to engage this pathway, contributing to the establishment and maintenance of the SASP ^29,38^. Whether similar mechanisms operate in post-mitotic neurons, particularly in the context of aging and disease when cGAS is upregulated in neurons^39,40^, remains unclear.

Therapeutically, the contribution of senescence to disease has motivated the development of senolytic agents, such as dasatinib and quercetin, which selectively eliminate senescent cells ^41^. While these approaches have shown promise in preclinical models, their application to neurodegenerative disease faces significant challenges, including limited blood-brain barrier penetration and the potential consequences of neuronal loss ^42,43^. An alternative strategy is the use of senomorphic interventions that suppress senescence-associated phenotypes without inducing cell death. In this context, inhibition of LINE-1 represents an attractive target. Nucleoside reverse transcriptase inhibitors (nRTIs), widely used as antiviral therapies, have been shown to suppress LINE-1 activity and attenuate inflammation in aging and disease models ^28,29,44,45^. Intriguingly, epidemiological analyses suggest that chronic nRTI exposure may reduce the incidence of age-associated inflammatory conditions, raising the possibility of repurposing these agents for neurodegenerative disease ^46^.

Here, we investigate the role of LINE-1 in neuronal senescence in Alzheimer’s disease (AD) and test the hypothesis that LINE-1 activity drives senescence-associated phenotypes through cytoplasmic DNA sensing pathways. Using induced neurons (iNs) directly converted from human fibroblasts that preserve donor-specific aging-associated molecular signatures, combined with long-read single-cell transcriptomics and spatial analyses of human brain tissue, we identify a subset of neurons with elevated LINE-1 activity that exhibit transcriptional signatures of senescence, neurodegeneration, and immune activation. We further demonstrate that LINE-1-derived cytoplasmic DNA activates cGAS-STING signaling in post-mitotic neurons and that inhibition of either LINE-1 or cytoplasmic DNA sensing suppresses these phenotypes. Together, our findings establish a mechanistic link between retrotransposon activity and neuronal senescence and identify the LINE-1/cGAS-STING axis as a tractable target for senomorphic therapy in neurodegenerative disease.

## Results

### Targeting LINE-1 reduces senescent phenotypes in AD neurons

Prior work demonstrated that senescent neurons can be selectively eliminated using the senolytic cocktail dasatinib and quercetin (DQ). However, quercetin was recently shown to have limited brain permeability in a clinical trial of DQ^43^, and the consequences of removing a substantial proportion of neurons from the AD brain remain unclear. An alternative approach is to preserve neuronal viability but suppress senescent phenotypes, an approach broadly termed senomorphic therapy. In multiple senescent cell types, inhibition of the retrotransposon long interspersed nuclear element 1 (LINE-1) prevents the development of senescent phenotypes through mechanisms including cytoplasmic DNAs, chromatin remodeling, and DNA damage responses ^33,47,48^. Given that neurons exhibit high levels of LINE-1 expression ^49^, we investigated whether inhibition of the LINE-1 life cycle with the nucleoside reverse transcriptase inhibitor (nRTI) 3TC could suppress senescence in AD neurons .

To test this, we prepared flow cytometry purified iNs from a cohort of AD and healthy age matched control patients. iNs retain the molecular, epigenetic, and proteomic signatures of aging that are otherwise lost in traditional neuron reprogramming strategies involving an induced pluripotent stem cell (iPSC) intermediate, and have been widely used model neuronal senescent phenotypes in vitro ^9,15,25^. Treatment with 3TC resulted in a dose-dependent reduction in p16INK4a, a canonical marker of senescence, in AD iNs but not control iNs (Fig 1A). At a concentration of 1 uM, 3TC reduced p16 expression in AD neurons to levels comparable to those observed in healthy controls (Fig 1B). This effect was neuron specific, as 3TC treatment of the corresponding parent fibroblast cultures did not alter p16 expression (Sup Fig 1 A). These findings were validated by immunohistochemistry for p16 following treatment with either 3TC and or a triple antisense oligonucleotide (ASO) cocktail targeting active human specific (L1Hs) elements (Sup Fig 1B), which we validated could significantly decrease active L1Hs expression (Sup Fig 1C).

**Figure 1:**
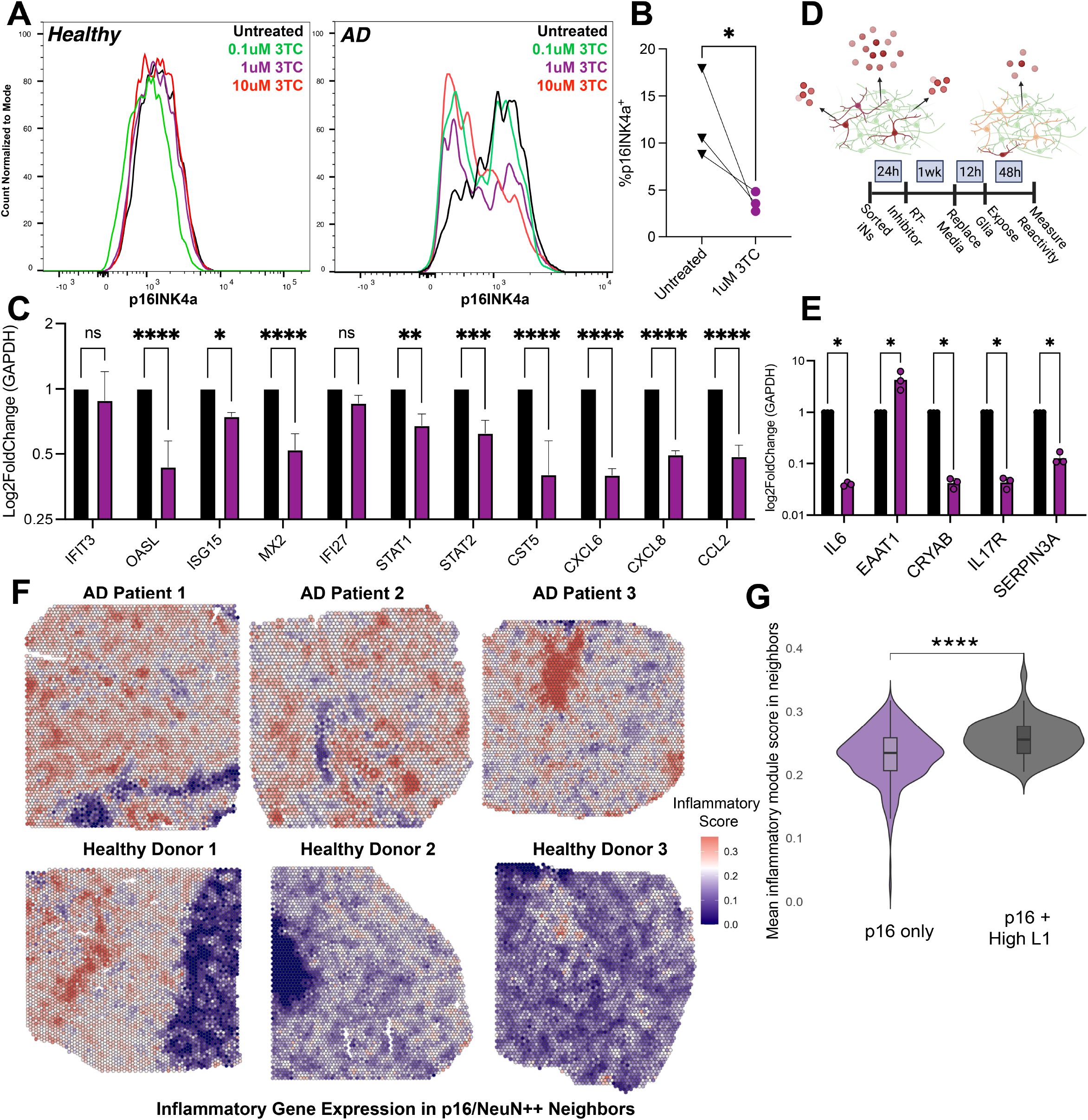
LINE-1 inhibition suppresses neuronal senescence and the SASP in Alzheimer’s disease iNs. (A) Flow cytometry analysis of p16 expression in induced neurons (iNs) derived from Alzheimer’s disease (AD) and age-matched control donors following treatment with increasing concentrations of the nucleoside reverse transcriptase inhibitor (nRTI) lamivudine (3TC), demonstrating a dose-dependent reduction in p16 in AD iNs. (B) Quantification of p16 expression showing that treatment with 1 ìM 3TC reduces p16 levels in AD iNs to those observed in healthy controls. * p < .05, paired t-test. (C) qPCR analysis of senescence-associated secretory phenotype (SASP) and interferon-stimulated genes (ISGs) in AD iNs ± 3TC treatment, showing significant downregulation upon LINE-1 inhibition. .** p < .01, *** p < .001, **** p < .0001, 2 way ANOVA. (D) Schematic of conditioned media (CM) experimental design. Neurons are senescent (red), healthy (green), or SASP suppressed (yellow). (E) Expression of reactive astrogliosis markers in astrocytes following CM exposure, demonstrating reduced astrocyte activation in response to CM from 3TC-treated AD iNs. 2 way ANOVA. (F) Spatial transcriptomic mapping of human middle temporal gyrus from AD and control donors. Cells neighboring p16+/L1:high neurons are colored by inflammatory gene module score. (G) Quantification of inflammatory gene expression in cells adjacent to p16+/L1:high versus p16+/L1:low neurons, showing increased inflammation in LINE-1 high neighborhoods. Wilcoxon signed rank test.

**Figure 2:**
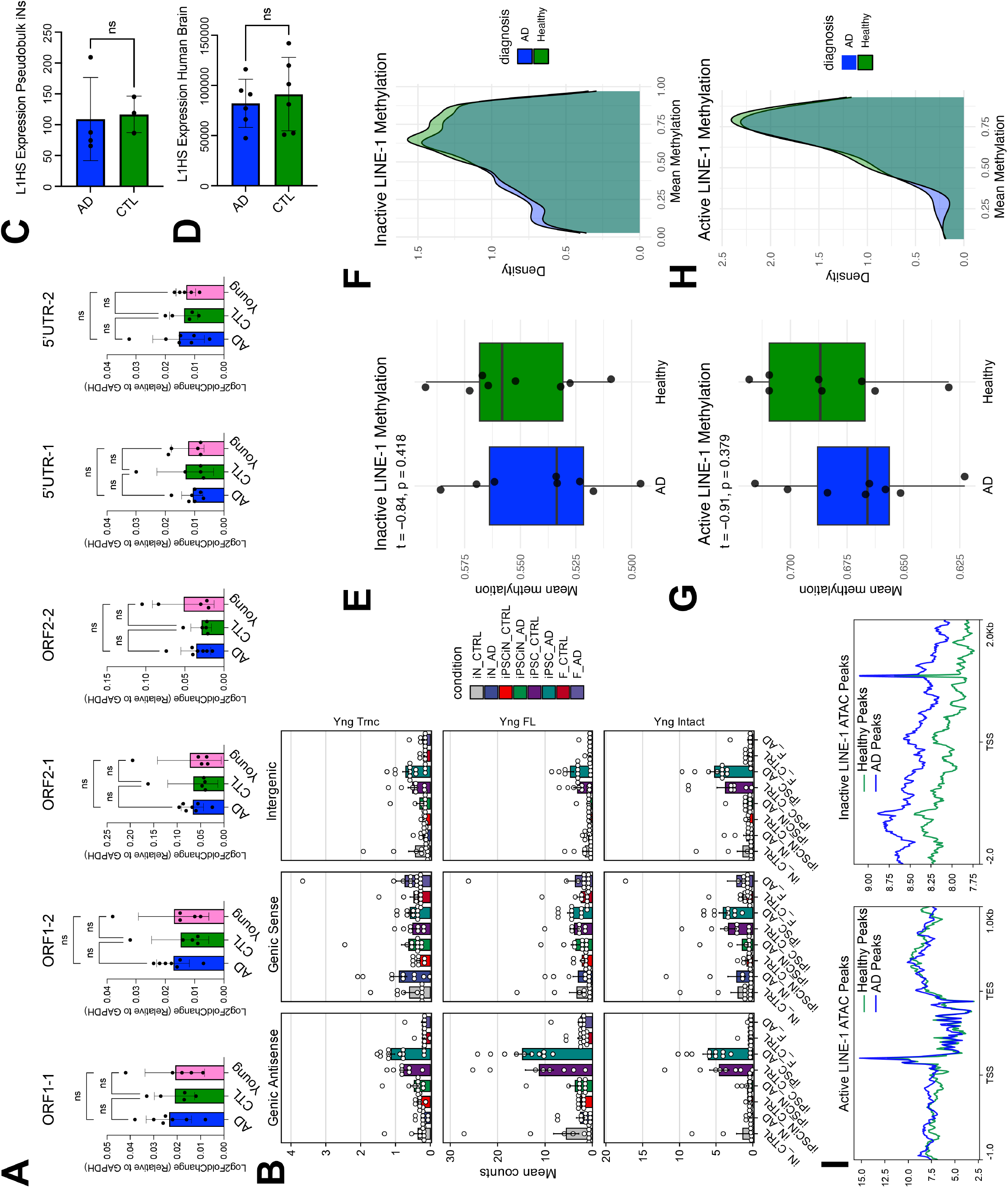
LINE-1 expression and epigenetic regulation are not globally altered in AD neurons. (A) qPCR analysis using L1Hs-targeted primer sets spanning ORF1, ORF2, and 5′UTR regions reveals no significant differences in active LINE-1 expression across AD, age-matched control, and young donor iNs. 2 way ANOVA. (B) Quantification of LINE-1 expression from short-read RNA-seq in iNs, fibroblasts, iPSCs, and iPSC-derived neurons. (C) L1Hs expression in bulk long-read RNA sequencing of postmortem human brain tissue in AD and control samples. Likelihood ratio test. (D) Pseudobulk analysis of long-read single-cell RNA-seq data confirming no significant differences in total L1Hs expression between AD and control neurons. Likelihood ratio test. (E–F) DNA methylation levels at CpG sites within inactive LINE-1 elements across AD and control samples. Un-paired t-test. (G–H) DNA methylation levels at CpG sites within active LINE-1 elements across AD and control samples. Un-paired t-test. (I) ATAC-seq chromatin accessibility profiles at LINE-1 loci.

Because reduced p16 expression is typically associated with loss of a senescence associated secretory phenotype (SASP), we next assess expression of a panel of SASP and interferon stimulated genes (ISGs). 3TC treatment significantly reduced expression of both SASP and ISG transcripts in AD iNs (Fig 1C). Given that paracrine signaling from senescent cells is thought to drive much of the deleterious tissue environment in aging, we examined whether 3TC treatment modulates the response of healthy human astrocytes to conditioned media from AD neurons (Fig 1D). Consistent with the observed reduction in p16, conditioned media from 3TC-treated AD iNs induced lower expression of reactive astrogliosis markers compared to untreated controls, indicating effective suppression of the SASP (Fig 1E).

To determine whether similar paracrine interactions could occur in human tissue, we analyzed a spatial transcriptomic dataset from the middle temporal gyrus of AD and healthy control donors^50^. Senescent neurons were identified based on co-expression of p16 and NeuN, and cells were stratified according to the abundance of reads mapping to LINE-1-containing sequences. We observed a significant increase in inflammatory gene module expression in cell neighboring p16+LINE-1:high neurons (top decile) compared to those adjacent to p16/LINE-1:low neurons (Fig1 F-G). Furthermore, AD samples exhibited a higher frequency of such p16/LINE-1:high neighborhoods relative to healthy controls (Sup Fig1 D). When comparing gene sets enriched in p16⁺/LINE-1⁺ neighborhoods, a notable finding was the increased expression of MHC-I in neighboring cells within AD samples, an effect not observed in healthy controls (Sup Fig1 E).

Together, these findings demonstrate that inhibition of LINE-1 activity suppresses neuronal senescence in AD iNs and support a model in which LINE-1 drives a pro-inflammatory, paracrine SASP both in vitro and in the human AD brain, accompanied by localized induction of inflammatory expression in neighboring cells.

### LINE-1 Expression Dynamics in Human Neurons

Next, we sought to define the mechanism by which LINE-1 inhibition modulates senescent phenotypes in AD iNs. Prior studies of LINE-1 expression in the human brain have yielded conflicting results, with some reports indicating increased expression in AD and others observing no significant differences ^25,51–54^. However, the sensitivity of AD iNs to both 3TC and ASO-mediated LINE-1 inhibition treatment raised the possibility that elevated LINE-1 expression could underlie the observed reductions in p16 and SASP factors.

A major challenge in studying LINE-1 biology is the high abundance of LINE-1 sequences in the human genome (>20%) and the difficulty in distinguishing intact, full length elements, which we refer to as “active” from evolutionarily degraded or truncated “passenger” copies, which we refer to as “inactive” ^55,56^. To address this, we developed and validated a set of six qPCR primer pairs spanning key domains of LINE-1, including ORF1, ORF2, and the 5′UTR, enabling preferential detection of intact, potentially active elements. We applied this panel to iNs derived from AD, age-matched healthy controls, and young donors to assess changes in active LINE-1 expression across aging and disease. Surprisingly, we detected no significant differences in LINE-1 expression between AD, healthy, or young neurons (Fig2 A).

To further interrogate LINE-1 expression, we leveraged a previously generated Illumina short-read RNAseq dataset from an expanded cohort, including iNs, fibroblasts, iPSCs, and iPSC-derived iNs lacking aging signatures ^20^. Using a specialized computational pipeline designed to distinguish active from inactive LINE-1 transcripts^57^, we again observed no significant differences in overall LINE-1 expression between AD and healthy samples (Fig2 B). Notably, undifferentiated iPSCs had the highest overall levels of LINE-1 expression in our dataset, consistent with the well-established upregulation of LINE-1 during early embryonic development.

Given the limitations of short read sequencing for resolving full-length active LINE-1 expression, we next analyzed long-read RNA sequencing datasets, which capture transcripts spanning kilobases and allow the resolution of the 5’ UTR, for identifying active elements, and the 3’ UTR and flanking regions that are specific to a unique genomic locus and identifying exact genomic origin. Applying a dedicated L1Hs-calling pipeline to published bulk long-read human brain data^58^, we again found no differences in LINE-1 abundance between AD and control samples (Fig2 C). Similar, pseudobulk analysis of long-read single-cell RNA-seq data (see Fig 3) revealed no significant differences in the expression of all L1Hs elements between AD and healthy iNs (Fig2 D).

**Figure 3:**
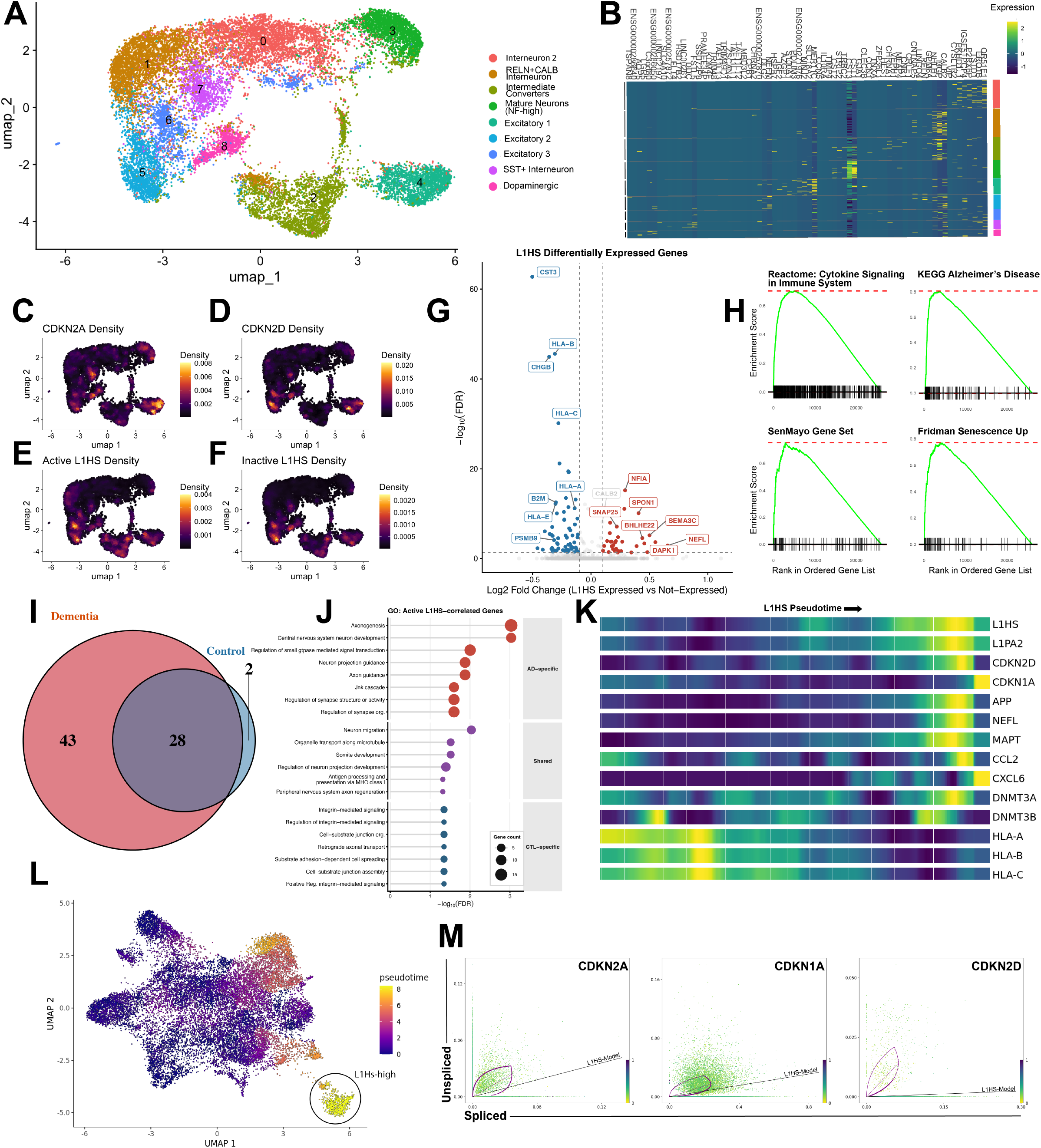
Single-cell long-read sequencing links LINE-1 activity to neuronal senescence trajectories. (A) UMAP clustering of iN single-cell long-read RNA-seq data. (B) Heatmap of top marker genes defining neuronal clusters. (C-D) Distribution of senescence marker expression p16 and p19 (E-F) Distribution of inactive and active L1 elements across clusters. (G) Volcano plot of differentially expressed genes between L1Hs+ and L1Hs- cells. (H) Gene set enrichment analysis (GSEA) of genes upregulated in L1Hs+ cells (I) Comparison of differential expression magnitude between dementia and control iNs. (J) Gene ontology analysis of genes correlated with L1Hs expression across patient conditions. (K) Pseudotime trajectory analysis ordering cells by increasing L1Hs expression. (L) Identification of a high-L1Hs state population along the trajectory. (M) RNA velocity phase portraits of senescence markers in reference to an L1Hs model, represented as the black line. Genes above this line have positive velocity, genes below have negative velocity.

LINE-1 activity has been implicated in modulating chromatin accessibility and DNA methylation ^59,60^. We therefore assessed whether epigenetic differences at LINE-1 loci distinguish AD from healthy neurons. Using the Illumina MethylEPIC platform, we analyzed CpG methylation at sites within inactive (Fig2 E-F) or active (Fig2 G-H) LINE-1 elements^20^. Active LINE-1 elements exhibited higher methylation levels than inactive elements, consistent with their repression by host defense mechanisms. While healthy samples showed a modest increase in methylation at both active and inactive LINE-1 CpGs, these differences were not significant.

We next re-analyzed a previously published ATAC-seq data from AD and healthy iNs to examine chromatin accessibility at LINE-1 loci ^20^. Using deepTools to generate aggregate accessibility profiles, we found that chromatin accessibility at active LINE-1 elements was comparable between AD and healthy neurons, consistent with their similar expression levels. In contrast, inactive LINE-1 regions exhibited significantly increased accessibility in AD samples (Fig2 I). However, given that inactive LINE-1 elements comprise a substantial fraction of the genome (∼20%), this effect likely reflects a broader pattern of chromatin relaxation rather than LINE-1-specific dysregulation. Indeed, we have previously reported widespread chromatin decompaction and dedifferentiation as hallmarks of the AD iN epigenome ^20,61^.

Taken together, these results indicate that LINE-1 is not broadly dysregulated at the level of expression or canonical epigenetic regulation in AD neurons in a bulk population, suggesting that the effects of LINE-1 inhibition are not driven by global changes in LINE-1 abundance, potentially reflecting heterogeneity with persistent alterations in a subset of individual cells.

### Single-cell trajectories link LINE-1 expression to neuronal senescence

As LINE-1 intervention altered senescence-associated phenotypes, and senescent cells represent a small fraction of the overall population, we hypothesized that bulk analyses may obscure transcriptional and epigenetic changes occurring within this minority. To address this limitation, and to distinguish full-length, potentially active LINE-1 elements from truncated copies, we employed a long-read single-cell approach using PacBio Kinnex RNA-seq.

We profiled 19,301 cells from 9 individuals (4 AD, 3 control, 2 AD Related-Dementia (ADRD)) using single-cell long-read RNA sequencing (PacBio MAS-seq), generating 83,674,058 full-length cDNA reads (S-reads) from 5,636,423 HiFi reads. After cell barcode assignment and quality filtering, we retained a mean of ∼1,918 reads per cell. To resolve the challenge of multi-mapping reads arising from repetitive transposable elements, we applied an expectation maximization algorithm that accurately redistributes counts from multi-mapping reads to distinct TE subfamilies ^62^. This approach enabled the generation of a unified single cell dataset integrating protein coding gene expression with LINE-1 derived counts, allowing joint analysis of canonical transcriptional programs and LINE-1 activity within the same cells. Our pipeline demonstrated high specificity, with only a small fraction of inactive LINE-1 elements misclassified as active which were excluded from downstream analyses (Sup Fig3 A).

Unsupervised clustering of transcriptomic profiles identified multiple neuronal subtypes, as well as a population of fibroblast-like undifferentiated cells which were removed for downstream analyses (Sup Fig3 B). Clustering of the remaining cells revealed distinct iN populations corresponding largely to cortical neuron subtypes, including interneurons, RELN+CALB interneurons, general excitatory, and a minority of cholinergic and dopaminergic neurons (Fig3 A-B). These subtypes are in agreement with other NGN2-ASCL1 transdifferentiation reports ^63–65^. We also observed a population of intermediate converters that had both neuronal and myogenic markers, as has also been observed in iN single cell experiments.

Senescent neurons, defined by expression of canonical markers including p16 (CDKN2A) and p19 (CDKN2D) comprised a minority population distributed across multiple excitatory neuronal subtypes (Fig3 C-D). Our pipeline further enabled discrimination of reads derived from active versus inactive LINE-1 elements, revealing that LINE-1 expression is broadly detectable across iNs (Fig3 E-F). Notably, we observed that mature excitatory neurons had the highest senescence score, concordant with previous *in vitro* and post-mortem studies (Sup Fig 3C) ^9,66,67^.

To directly assess the transcriptional impact of LINE-1 activity, we stratified cells based on expression of full-length active human specific L1Hs and performed differential expression analysis between L1Hs-positive and L1Hs-negative cells (Fig3 G). Genes enriched in L1Hs-expressing cells were associated with neurodegeneration, immune signaling, and senescence pathways (Fig 3H). Conversely, L1Hs-positive cells exhibited downregulation of MHC-I genes (HLA-A, HLA-B, HLA-C), oxidative phosphorylation and electron transport, and proteasome assembly genes (Fig3 G, Sup Fig3 D). These transcriptional changes were more pronounced in iNs derived from dementia patients, which exhibited a greater number of differentially expressed genes compared to controls, suggesting an amplified transcriptional response to LINE-1 activity in disease contexts (Fig3 I, Sup Fig 3 E). Genes correlated with L1Hs expression in both AD and control neurons were enriched for more general neuronal functions including neuron migration, organelle transport along microtubules, and regulation of neuron projection development, as well as loss of antigen processing and presentation via MHC class I (Fig3 J).

Given the sensitivity of senescent phenotypes in iNs to LINE-1-modulating interventions, we next investigated the relationship between LINE-1 activity and cellular senescence at single-cell resolution. Cells were ordered along a pseudotime trajectory corresponding to increasing L1Hs expression, and genes varying along this continuum were identified using Monocle3 (Fig3 K-L)^68^. This analysis revealed a distinct population of cells occupying a high L1Hs-expression state (Fig 3L). Genes positively correlated with increasing L1Hs pseudotime included canonical AD associated markers (MAPT, APP, NEFL), as well as interferon-responsive genes and loss of MHC-I expression, consistent with our differential expression results (Fig3 G,K). Senescence markers (CDKN2A/p16, CDKN1A/p21) and SASP-associated factors (e.g., CCL2, CXCL6) also increased along this trajectory (Fig3 K). Interestingly, expression of DNA methyltransferases (DNMT3A and DNMT3B) was elevated with increasing LINE-1 expression, potentially reflecting a compensatory response aimed at re-establishing epigenetic repression.

Although LINE-1 activity has been linked to senescence in multiple systems, the temporal relationship between these processes remains unclear. To determine whether LINE-1 activation precedes or follows the onset of senescence-associated transcriptional programs, we performed RNA velocity analysis using the scVelo dynamical model ^69^. Cells were ordered along a latent time axis aligned with increasing LINE-1 expression. Analysis of gene-specific phase portraits revealed that senescence markers (p16, p21, p19) follow the trajectory of LINE-1 activation, indicating that senescence-associated transcriptional programs emerge downstream of LINE-1 expression (Fig3 M). These findings support a model in which aberrant LINE-1 activity contributes to the induction of neuronal senescence, rather than arising as a secondary consequence of it. Together, our long-read single-cell approach overcomes key limitations in resolving LINE-1 activity and detecting rare senescent populations, revealing a robust and temporally structured relationship between LINE-1 activation and neuronal senescence that is not apparent from bulk analyses.

### LINE-1 derived cytoplasmic DNA accumulates in post-mitotic neurons

Having established a transcriptional link between LINE-1 expression and activation of SASP programs, we next sought to define the mechanism by which LINE-1 activity promotes a neuronal SASP. LINE-1 has been reported to induce sterile inflammatory responses in senescent cells through accumulation of cytoplasmic DNA, which activates the cGAS–STING pathway and downstream interferon signaling.

To test for the presence of cytoplasmic DNA in our AD iN system, we employed an EdU labeling strategy coupled with click chemistry to mark newly synthesized DNA over a 48 hour period (Fig4 A). Notably, the majority of newly synthesized DNA was mitochondrial in origin (Fig4 B, Sup Fig4 A). To isolate non-mitochondrial signal, we applied a TOM20-based mask to exclude mitochondrial associated EdU signal. We observed a significant reduction in TOM20-negative, EdU-positive cytoplasmic puncta in iNs treated with two nRTIs, as well as a LINE-1-targeting ASO (Fig4 C). These results suggest that a subset of cytoplasmic DNA in iNs is dependent on LINE-1 activity. Importantly, overall TOM20 signal was unchanged between nRTI and untreated groups, arguing against differences in mitochondrial content as a confounding factor (Sup Fig4 B).

While these findings support a link between LINE-1 activity and cytoplasmic DNA accumulation, the EdU based approach does not distinguish the origin or sequence identity of the labeled DNA. To directly visualize LINE-1-derived DNA, we developed a CRISPR-based SunTag imaging strategy (Fig4 D)^70^. In this system, a nuclease-dead Cas9 (dCas9) fused to a nuclear export sequence and tandem GCN4 peptide repeats was stably expressed, enabling signal amplification through recruitment of an scFv:GCN4:sfGFP fusion protein. A guide RNA targeting the 5′ UTR of the human-specific LINE-1 element (L1Hs) was delivered via a separate lentivirus, allowing sequence-specific labeling of LINE-1 DNA in live cells.

To validate the specificity of this system, cells were transfected with a plasmid containing LINE-1 sequence or a control construct (Sup Fig4 C). Robust cytoplasmic fluorescence was observed selectively in LINE-1-transfected cells, confirming target specificity. We next assessed the sensitivity of L1Hs::SunTag to the effect of LINE-1-targeting drugs on endogenous LINE-1 using a longitudinal live imaging system for measuring GFP signal. Following an 8-hour pretreatment, ASO and nRTI conditions (3TC and FTC) all resulted in a reduction in L1Hs::SunTag signal intensity relative to untreated controls (Fig4 E). These findings were independently validated by high-resolution microscopy (Fig 4F-G). We then examined the subcellular localization of LINE-1 DNA across neuronal conversion. Interestingly, we observed a marked increase in cytoplasmic LINE-1 signal by day 14 of neuronal differentiation, whereas signal in parental fibroblasts was predominantly nuclear (Fig4 H). While the absence of cytoplasmic signal in fibroblasts may reflect nuclear sequestration of the dCas9 complex during cell division, it is also consistent with the low baseline levels of LINE-1 expression reported in fibroblasts^71^. These observations raise the possibility that post-mitotic neuronal states permit greater accumulation of cytoplasmic LINE-1 DNA. Finally, we investigated the relationship between LINE-1 DNA accumulation and cellular senescence. High-resolution imaging revealed that p16-positive cells exhibited significantly higher L1Hs::SunTag signal intensity compared to p16-negative cells, indicating an association between cytoplasmic LINE-1 DNA and senescent states (Fig4 I). Together, these results demonstrate that aged AD neurons accumulate LINE-1-derived DNA within the cytoplasm, supporting a model in which cytoplasmic DNA sensing pathways contribute to the induction of a neuronal SASP downstream of LINE-1 activity.

**Figure 4:**
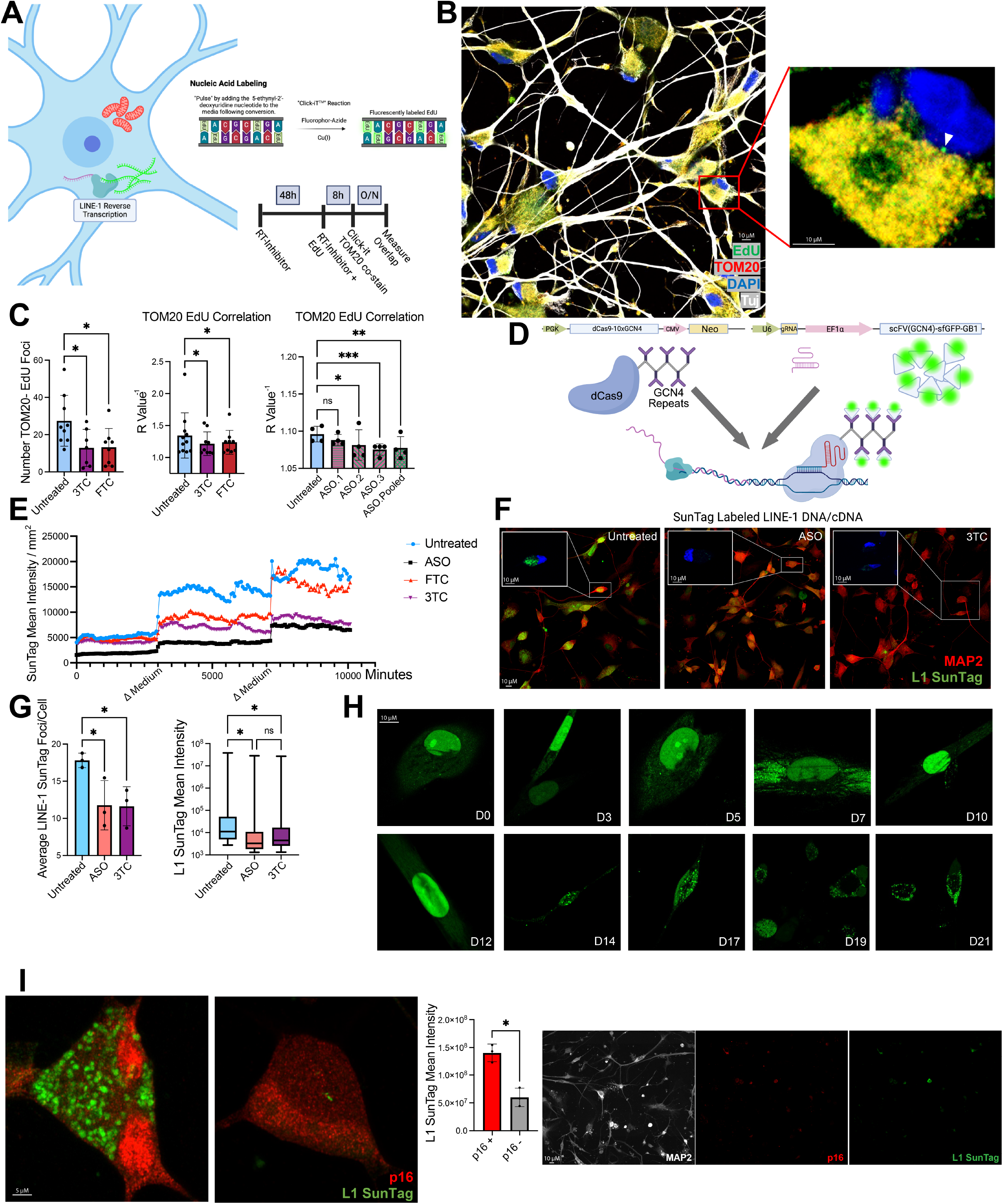
LINE-1-derived cytoplasmic DNA accumulates in post-mitotic neurons. (A) Schematic of EdU incorporation assay used to label newly synthesized DNA in iNs. (B) Representative images showing EdU labeling, arrowhead referencing a TOM20- EdU+ signal. (C) Quantification of TOM20-negative EdU-positive cytoplasmic signal following treatment with nRTIs or LINE-1-targeting ASOs. * p < .05, ** p < .01, *** p < .001. Paired t-test. (D) Schematic of CRISPR-based SunTag imaging system for visualization of LINE-1 DNA using dCas9-GCN4 and scFv-GFP amplification targeting the L1Hs 5′UTR. (E) Quantification of longitudinal L1Hs::SunTag signal intensity following treatment with ASOs and nRTIs (3TC, FTC).. (F) High-resolution microscopy images and of cytoplasmic LINE-1 DNA upon pharmacological inhibition. (G) quantification of (F). Paired t-test. (H) Subcellular localization of LINE-1 DNA during neuronal conversion, showing increased cytoplasmic accumulation in post-mitotic iNs (d21) compared to parental fibroblasts (d0). (I) Quantification of L1Hs::SunTag signal in p16+ versus p16- cells, demonstrating enrichment of cytoplasmic LINE-1 DNA in senescent neurons. Paired t-test.

**Figure 5.**
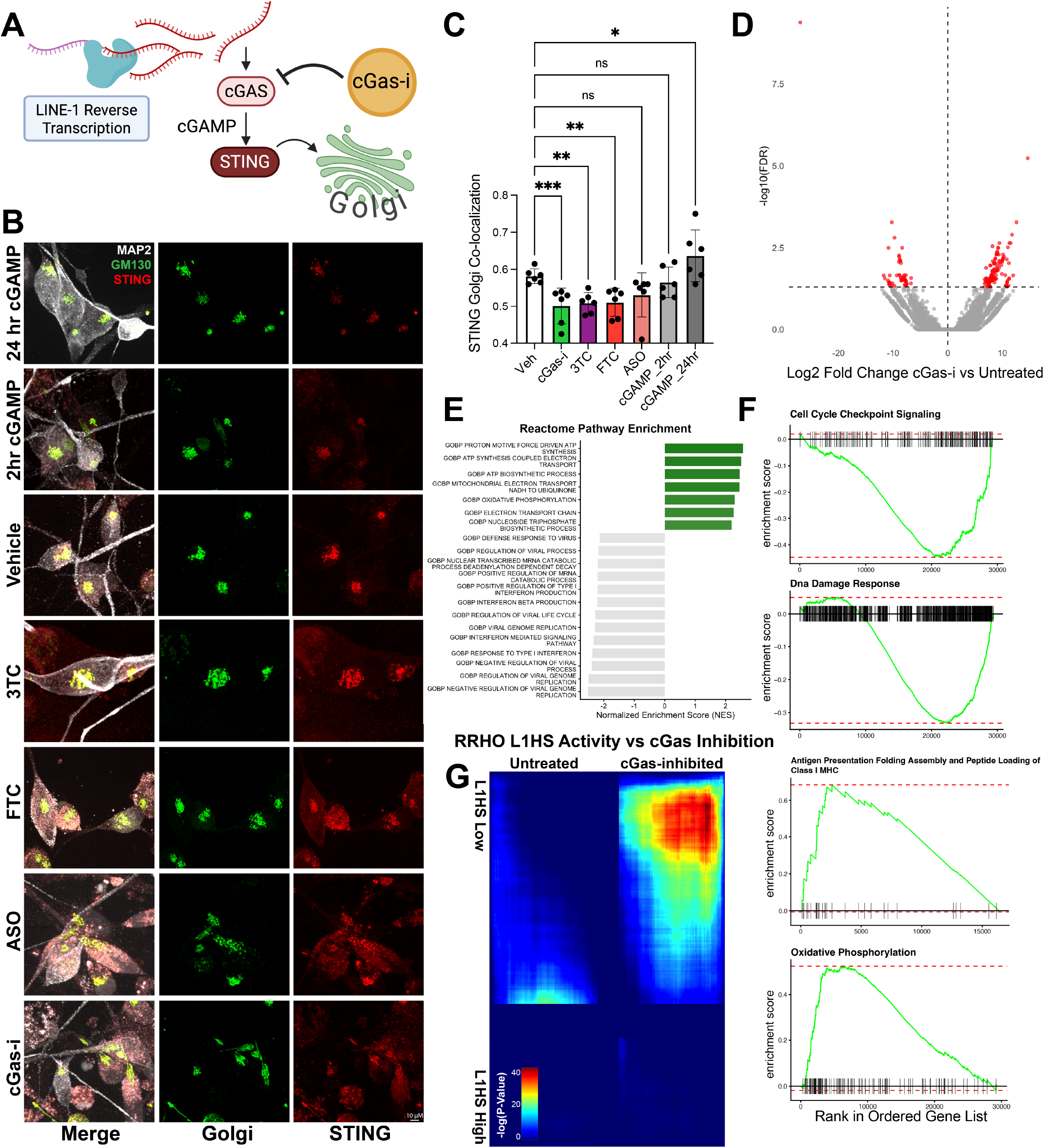
Inhibition of cGAS–STING signaling phenocopies LINE-1 suppression. (A) Schematic illustrating STING translocation to the Golgi as a readout of cGAS–STING activation. (B) Immunofluorescence images of STING and Golgi (GM130) localization in iNs following treatment with nRTIs, LINE-1 ASOs, a cGAS inhibitor (cGAS-i), or cGAS activator (cGAMP). (C) Quantification of STING and Golgi colocalization following treatment with nRTIs, LINE-1 ASOs, or a cGAS inhibitor (cGAS-i).* p< .05, ** p < .01, *** p < .001. 2 way ANOVA (D) Volcano plot of differentially expressed genes following cGAS inhibition in AD iNs compared to vehicle-treated controls. (E) GSEA showing enrichment of interferon and viral response pathways in untreated cells and oxidative phosphorylation and biosynthetic pathways upon cGAS inhibition. (F) Expression of dedifferentiation and MHC-I–related gene programs in untreated versus cGAS-inhibited cells. FDR q values < .001 for all. (G) Rank–rank hypergeometric overlap (RRHO) analysis comparing cGAS inhibition signatures with LINE-1 pseudotime associated genes, higher ranked list overlap is represented by warmer colors demonstrating strong concordance between low LINE-1 activity and cGAS inhibition

### Inhibition of cytoplasmic DNA sensing resolves AD related stress

Cytoplasmic, membrane unbound DNA is a well-established trigger of innate immune signaling in eukaryotic cells, eliciting a “non-self” DNA response across diverse contexts ^72–74^. The presence of LINE-1 DNA in the cytoplasm of AD iNs prompted us to test whether inhibition of cytoplasmic DNA sensing could phenocopy the effects of LINE-1-targeting interventions. To this end, we treated cells with a potent cGAS inhibitor (cGAS-i), alongside two nucleoside reverse transcriptase inhibitors (nRTIs), LINE-1-targeting ASOs, and cGAMP as a positive control. Activation of the cGAS–STING pathway was assessed by measuring STING translocation to the Golgi apparatus (Fig5 A-B). We observed that both nRTIs and cGAS inhibition markedly reduced STING translocation, with LINE-1 ASOs producing a more modest effect (Fig5 C). These results indicate that LINE-1 activity contributes to activation of the cGAS–STING pathway in AD neurons.

To further characterize the transcriptional consequences of cGAS inhibition, we performed bulk RNA-seq following one week of cGAS-i treatment compared to vehicle controls. Differential expression analysis identified 130 upregulated and 52 downregulated genes in response to cGAS inhibition (Fig5 D). Gene set enrichment analysis revealed that untreated cells exhibited increased expression of pathways associated with viral life cycle processes and interferon signaling, whereas cGAS-inhibited cells showed enrichment for oxidative phosphorylation and nucleoside biosynthesis pathways (Fig5 E). Notably, untreated cells also displayed transcriptional features consistent with dedifferentiation, alongside increased expression of MHC-I genes (Fig5 F). To integrate these findings with our single-cell analyses, we performed rank–rank hypergeometric overlap (RRHO) between the cGAS inhibition dataset and genes associated with LINE-1 pseudotime. This analysis revealed that cells with low L1Hs expression exhibited strong transcriptional similarity to cGAS-inhibited cells, suggesting that diminished cytoplasmic DNA sensing recapitulates a state of reduced LINE-1 activity (Fig5 G).

Collectively, these results support a model in which LINE-1-derived cytoplasmic DNA activates cGAS-STING signaling, thereby linking LINE-1 activity to neuronal senescence and inflammation in Alzheimer’s disease. Pharmacological inhibition of either LINE-1 activity or cytoplasmic DNA sensing attenuates this pathway, highlighting both as potential senotherapeutic targets. These findings provide a framework for the development of brain-penetrant LINE-1-directed therapies for neurodegenerative disease.

## Discussion

Cellular senescence has emerged as a central contributor to aging and neurodegenerative disease, yet therapeutic strategies to target senescent cells in the brain remain limited. While senolytic approaches such as dasatinib and quercetin (DQ) can effectively eliminate senescent neurons *in vitro*, their translational potential is constrained by poor blood-brain-barrier penetration. Here, we instead pursued a senomorphic strategy aimed at suppressing senescence phenotypes while preserving neuronal viability. Our findings identify LINE-1 as a key regulator of neuronal senescence in Alzheimer’s disease (AD) and demonstrate that pharmacological or genetic inhibition of LINE-1 activity attenuates both cell-intrinsic and paracrine features of the senescence-associated secretory phenotype (SASP).

We show that inhibition of LINE-1 activity, using either nucleoside reverse transcriptase inhibitors (nRTIs) or antisense oligonucleotides, reduces canonical senescence markers, including p16, and suppresses SASP and interferon-stimulated gene expression in AD neurons. Conditioned media experiments demonstrate that this effect extends beyond cell-autonomous changes, reducing the ability of senescent neurons to induce reactive states in neighboring astrocytes. Consistent with these *in vitro* findings, spatial transcriptomic analyses of human AD brain tissue reveal that neurons with high LINE-1 expression and senescence markers are associated with localized inflammatory niches and altered MHC-I expression in neighboring cells. These results support a model in which LINE-1 activity contributes to a pro-inflammatory, paracrine signaling environment in the AD brain. This is consistent with prior work implicating retrotransposon activation in sterile inflammation but extends these observations to post-mitotic human neurons and highlights LINE-1 as a therapeutically targetable node for modulating the neuronal SASP.

A striking finding from our study is the apparent disconnect between the functional impact of LINE-1 inhibition and the lack of detectable differences in LINE-1 expression at the bulk level between AD and control neurons. Across multiple orthogonal approaches including targeted qPCR, short-read RNA-seq, long-read bulk RNA-seq, DNA methylation analyses, and chromatin accessibility we observed no significant global changes in active LINE-1 abundance or canonical epigenetic regulation. This discrepancy highlights the challenge in studying repetitive elements in bulk measurements which can obscure rare but functionally significant cell states. Our long-read single-cell RNA sequencing approach resolves this limitation by enabling simultaneous detection of full-length, active LINE-1 transcripts and host gene expression within individual cells. Using this strategy, we identify a subset of neurons characterized by high LINE-1 activity that exhibit transcriptional signatures of neurodegeneration, immune activation, and senescence. These LINE-1 high cells are enriched in AD samples and display amplified senescence gene expression relative to controls, suggesting increased vulnerability or responsiveness in disease contexts. These findings support a model in which LINE-1 driven phenotypes are not governed by global upregulation, but rather by cellular heterogeneity and activation within a sub population of neurons. This may help reconcile conflicting reports in the literature regarding LINE-1 expression in AD and aging and underscores the importance of single-cell approaches for resolving transposable element biology in complex tissues.

The causal relationship between LINE-1 activation and senescence has remained unresolved, with prior studies suggesting that retrotransposon reactivation is a late consequence of chromatin breakdown in senescent cells or loss of L1 regulators^75,76^. In contrast, our single-cell trajectory and RNA velocity analyses indicate that LINE-1 activation precedes the induction of canonical senescence markers, including p16, p19, and p21, in neurons. Our results suggest aberrant LINE-1 activity contributes to the initiation of senescence-associated transcriptional programs, rather than arising solely as a downstream byproduct. One interpretation is that low-level or stochastic activation of LINE-1 in vulnerable neurons may act as an early trigger for stress responses that culminate in senescence. At the same time, our data do not exclude a feedback model in which senescence-associated chromatin remodeling further amplifies LINE-1 activity, reinforcing inflammatory signaling.

Mechanistically, our data support a model in which LINE-1 promotes neuronal senescence through the generation of cytoplasmic DNA that activates innate immune signaling pathways. Using both EdU labeling and a CRISPR-based SunTag imaging system, we demonstrate that LINE-1-derived DNA accumulates in the cytoplasm of post-mitotic neurons and is enriched in senescent cells. Pharmacological inhibition of LINE-1 reduces this cytoplasmic DNA pool, indicating that a subset of cytosolic DNA in neurons arises from active reverse transcription. Notably, our data do not exclude other sources of cytoplasmic DNA in neurons, however, only the LINE-1-dependent fraction is sensitive to reverse transcriptase inhibition and associated with senescent phenotypes, suggesting that LINE-1 sources of cytoplasmic DNA may have a particular impact on neuronal senescence. The enrichment of cytoplasmic LINE-1 DNA in post-mitotic neurons is particularly intriguing and may reflect cell-type specific differences in nuclear envelope dynamics, LINE-1 silencing pathways, or LINE-1 regulation through macromolecular condensates. These observations raise the possibility that neurons are uniquely susceptible to accumulation of retrotransposon-derived DNA, potentially contributing to their vulnerability in aging and neurodegeneration.

Consistent with the presence of cytoplasmic DNA, we find that inhibition of the cGAS–STING pathway phenocopies key aspects of LINE-1 inhibition, including reduced interferon signaling and altered metabolic and differentiation states. Transcriptomic analyses reveal that cGAS inhibition shifts neurons away from viral response and dedifferentiation programs toward oxidative phosphorylation and biosynthetic pathways, suggesting a broader restoration of cellular homeostasis. Integration of bulk and single-cell datasets further supports a close relationship between LINE-1 activity and cytoplasmic DNA sensing, as cells with low LINE-1 expression exhibit transcriptional similarity to cGAS-inhibited cells. Together, these findings place LINE-1 upstream of cGAS–STING activation in neurons and identify cytoplasmic DNA sensing as a key mediator of LINE-1-driven senescence phenotypes.

Our findings identify LINE-1 as a promising target for senomorphic therapies in neurodegenerative disease. Unlike senolytic approaches, which eliminate cells, inhibition of LINE-1 activity suppresses deleterious inflammatory phenotypes while preserving neuronal populations. The use of clinically approved nRTIs, such as lamivudine (3TC), provides a potential translational pathway, although optimization of brain penetration and specificity will be critical. More broadly, this work highlights transposable elements as active regulators of cellular state rather than passive genomic elements. Targeting LINE-1 or its downstream signaling pathways may therefore represent a generalizable strategy for modulating senescence and inflammation across tissues.

## Supplemental Figures

**Sup Figure 1.**
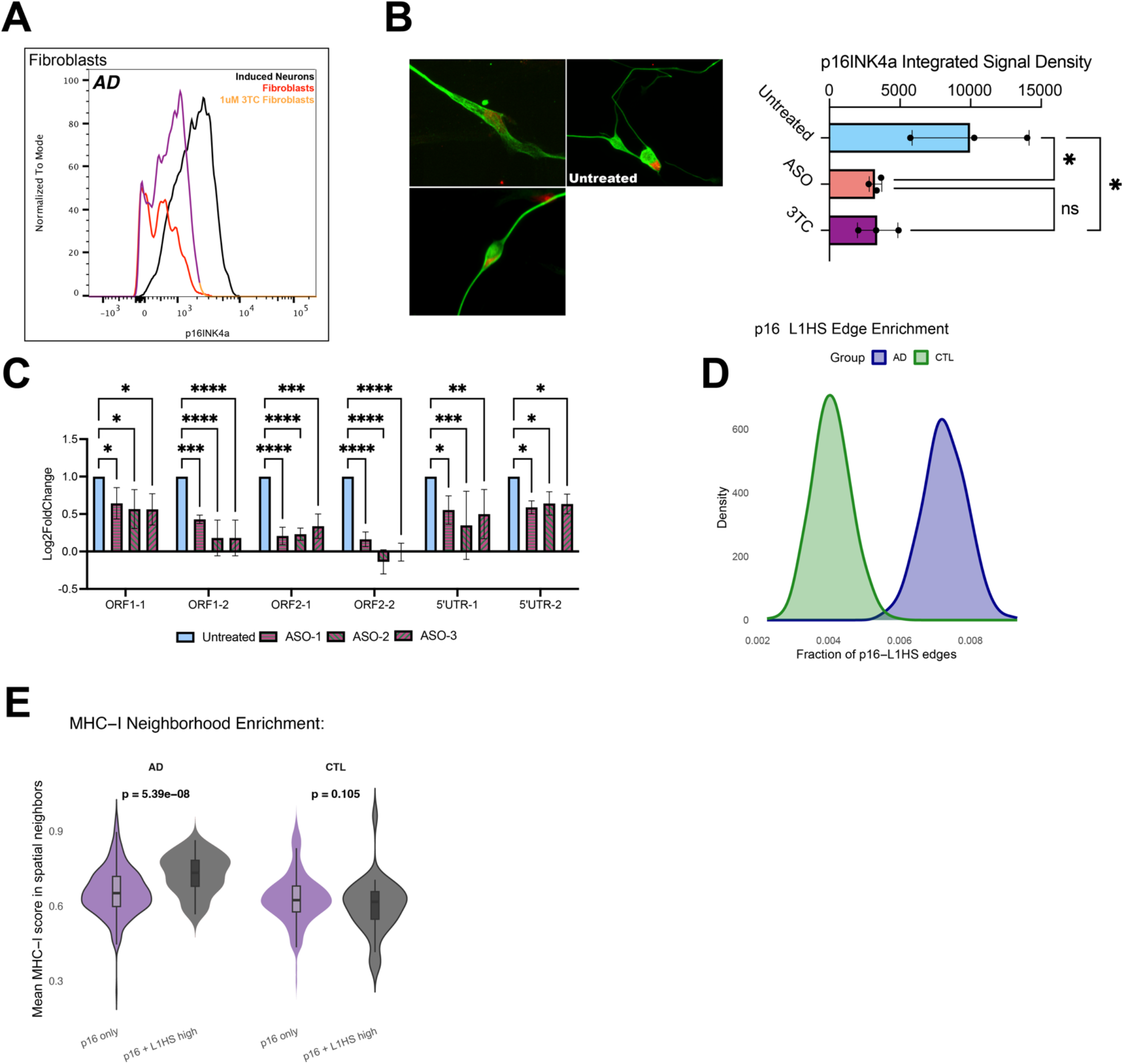
(A) Quantification of p16^INK4a^ expression in parental fibroblast cultures derived from Alzheimer’s disease (AD) and age-matched control donors following treatment with lamivudine (3TC). (B) Representative immunofluorescence images and quantification of p16 expression in AD iNs following treatment with lamivudine (3TC) or a triple antisense oligonucleotide (ASO) cocktail targeting active human-specific LINE-1 (L1Hs) elements. * p < .05, paired t-test. (C) Validation of LINE-1 ASO efficacy by qPCR analysis of active L1Hs expression following treatment with the triple ASO cocktail. ** p < .01, *** p < .001, **** p < .0001. 2 way ANOVA. (D) Quantification of spatial transcriptomic neighborhoods in human middle temporal gyrus samples showing increased abundance of p16⁺/LINE-1:high neuronal niches in AD relative to healthy control donors. (E) Gene expression analysis of neighboring cells highlighting increased MHC-I expression in AD-specific p16+/L1:high niches. Wilcoxon signed rank test.

**Sup Figure 3.**
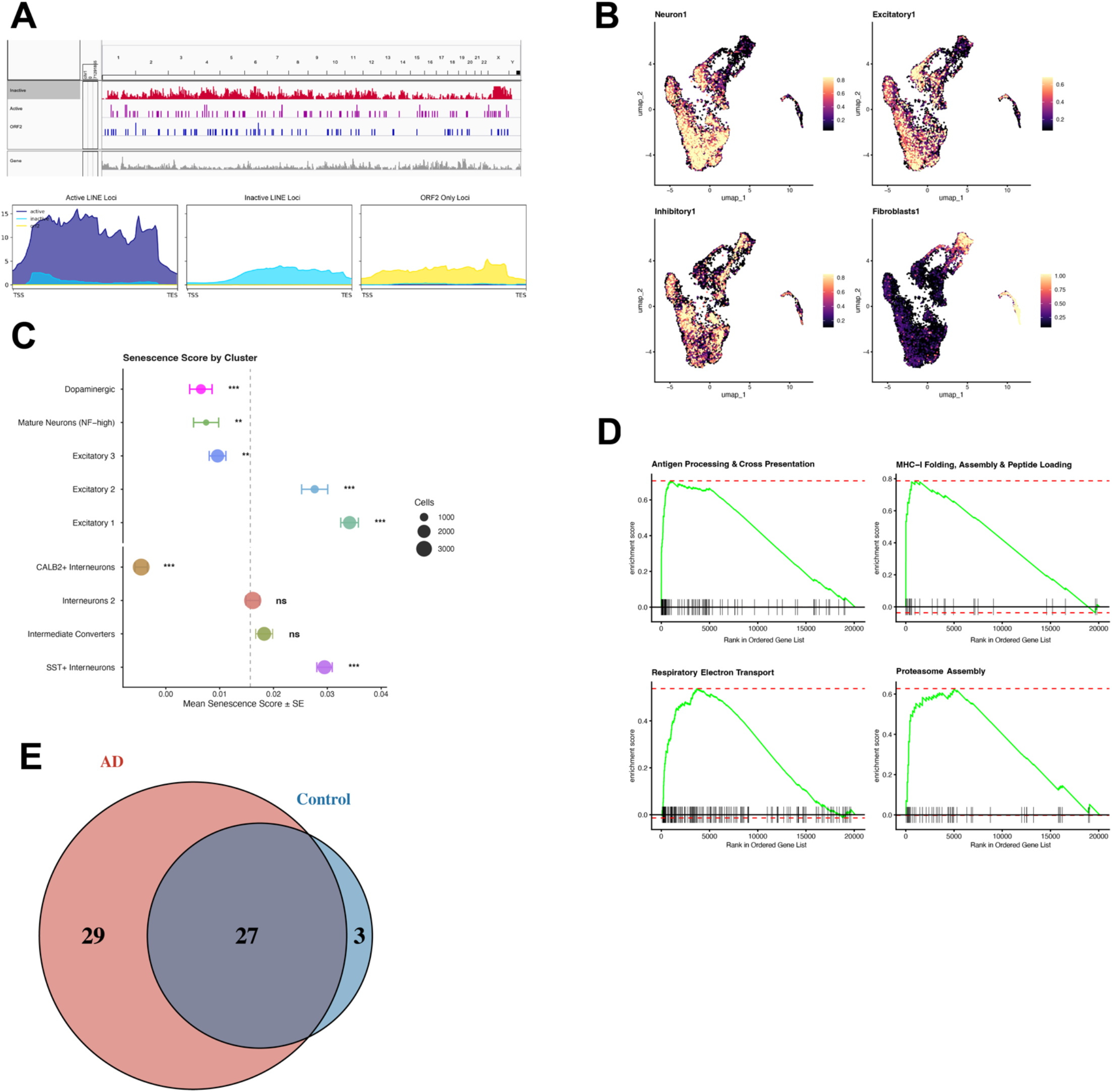
(A) Validation of the long-read LINE-1 quantification pipeline. Classification performance of active versus inactive LINE-1 (L1Hs) elements following application of an expectation–maximization–based read assignment strategy to resolve multimapping transposable element reads. (B) UMAP visualization and marker-based annotation of all sequenced cells prior to filtering, highlighting a population of fibroblast-like undifferentiated cells that were excluded from downstream neuronal analyses. (C) Senescence score distribution across neuronal subtypes identified by long-read single-cell RNA sequencing. (D) Gene set enrichment analysis (GSEA) of genes downregulated in L1Hs+ cells. FDR q values < .001 for all. (E) Comparison of differential expression magnitude between Alzheimer’s and control iNs.

**Sup Figure 4.**
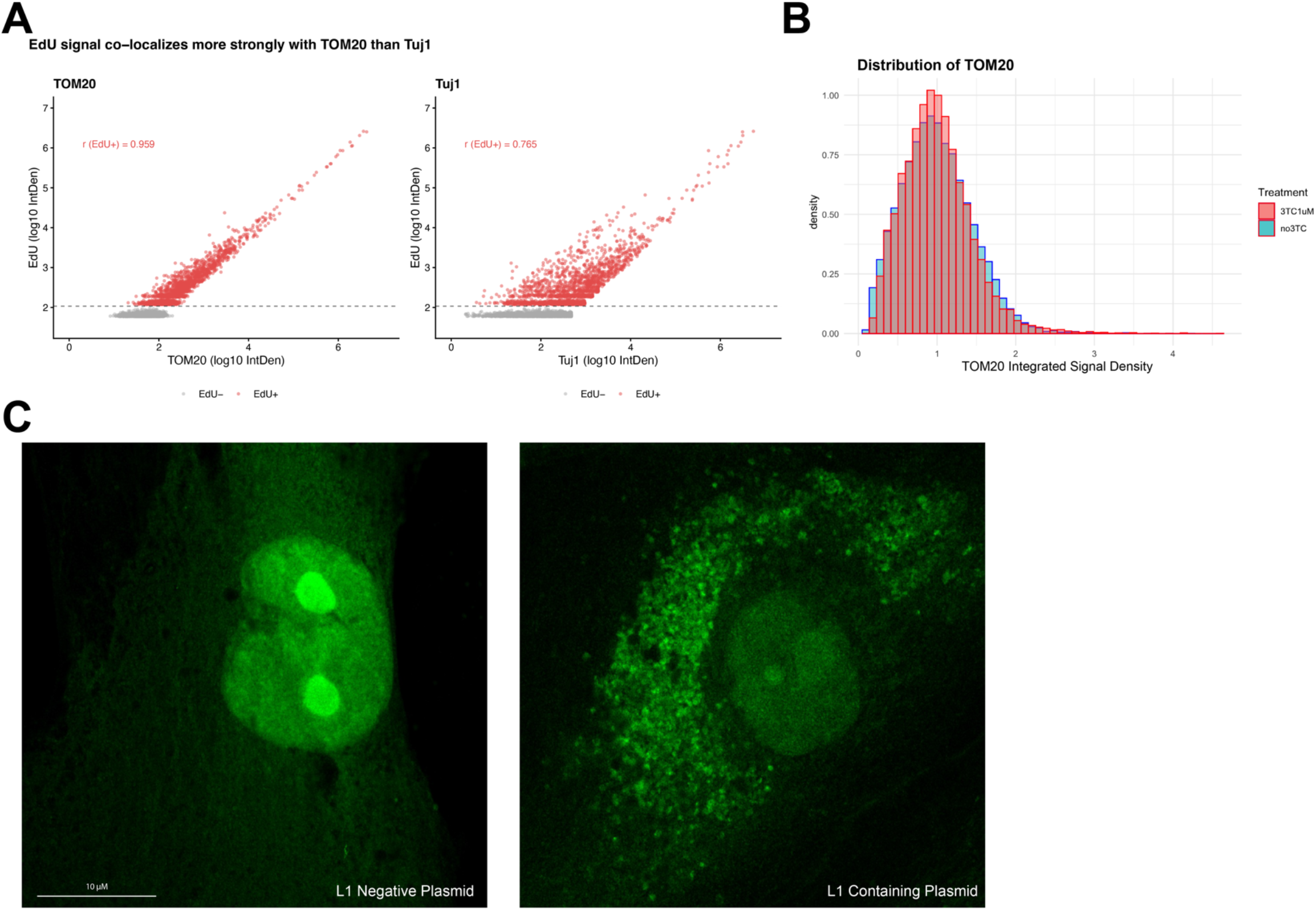
(A) Quantification of EdU incorporation in induced neurons (iNs), demonstrating that the majority of newly synthesized DNA signal colocalizes with the mitochondrial marker TOM20, consistent with predominantly mitochondrial DNA synthesis. (B) Quantification of TOM20 signal intensity following treatment with 3TC. No significant differences in TOM20 abundance were observed during treatment conditions. (C) Validation of the CRISPR-based L1Hs::SunTag imaging system. Cells expressing nuclease-dead Cas9 (dCas9)-SunTag and a guide RNA targeting the 5′ untranslated region (UTR) of active human-specific LINE-1 (L1Hs) elements were transfected with either a LINE-1-containing construct or control plasmid. Robust cytoplasmic fluorescence was selectively observed in LINE-1-transfected cells, confirming target specificity of the imaging strategy.

## ACKNOWLEDGEMENTS

We are grateful to all donors participating in this study. Special thanks to Gage lab members Meiyan Wang, Raffaella Lucciola, Jeff Jones, Elisa Gorostieta-Salas, Mary Lynne Gage, and Lisa Mitchell for scientific advice and editorial support.

This work was supported by the BrightFocus Foundation award A2025010F and we thank the Foundation Staff for their support and helpful discussions.

EDUC2-12611 CIRM Bridges to Stem Cell Research and Therapy Training Grant MiraCosta College, NSF ExLENT Award #<u>2322395</u>

This work was supported by Daniela Boassa, Chynna Bowman and Elsie Quansah from the Waitt Advanced Biophotonics Core Facility of the Salk Institute (RRID:SCR_014838) with funding from NIH-NCI CCSG P30 CA014195, NIH-NIA San Diego Nathan Shock Center P30 AG068635, The Henry L. Guenther Foundation and the Waitt Foundation

This work was supported by the Cell Technologies and Engineering Core Facility of the Salk Institute (RRID:SCR_014850) with funding from the NIH-NIA San Diego Nathan Shock Center P30 AG068635, the NIH-NIA Alzheimer’s Disease Research Center P30 AG062429, the AHA Allen Initiative, the California Institute for Regenerative Medicine and the Helmsley Charitable Trust.

This work was supported by Caz O’Connor, Mikayla Marrin, and Michelle Liem from the Flow Cytometry Core Facility of the Salk Institute (RRID:SCR_014839) with funding from NIH-NCI CCSG P30 CA014195, and Shared Instrumentation Grants S10-OD023689 (Aria Fusion cell sorter), and S10 OD034268 (Thermo Fisher Bigfoot).

We acknowledge support from P01 AG051449, NIH grant #R37 AG072502, the Dolby Family Fund, the Arnold and Mabel Beckman Foundation, The Freedom Together Foundation, AHA-Allen Initiative in Brain Health and Cognitive Impairment award made jointly through the American Heart Association and The Paul G. Allen Frontiers Group: 19PABH134610000.

We would like to thank Longevity Consortium (award # U19 AG023122 and award # U19AG023122-16), and the NIH-NIA grant P01 AG051449, P30 AG062429, and R37 AG072502.

We thank the Stem Cell Genomics Core at the Sanford Stem Cell Institute for providing sequencing services

## METHODS

### Direct Conversion of Human Fibroblasts into iNs

Primary human dermal fibroblasts from donors between 0 and 88 years of age were obtained from the Coriell Institute Cell Repository, the University Hospital in Erlangen and Shiley-Marcos Alzhiemer’s Disease Research Center (Table?). Protocols were previously approved by the Salk Institute Institutional Review Board and informed consent was obtained from all subjects. Fibroblasts were cultured in DMEM containing 15% tetracycline-free fetal bovine serum and 0.1% NEAA (Thermo Fisher Scientific), transduced with lentiviral particles for EtO and XTP-Ngn2:2A:Ascl1 (E+N2A), or the combined tetOn system cassette consisting of the rtTAAdv. [Clonech] driven by the UbC promoter, Ngn2:2A:Ascl1 under control of the TREtight promoter [Clontech], and a puromycin-resistance gene driven by the PGK promoter (UNA, Sup Fig2A) and expanded in the presence of puromycin (1 µg/ml; Sigma Aldrich) as ‘iN-ready’ fibroblast cell lines. Following at least three passages after viral transduction, ‘iN-ready’ fibroblasts were trypsinized and pooled into high densities (30,000 – 50,000 cells per cm2 ; appx. a 2:1 – 3:1 split from a confluent culture) and, after 24h, the medium was changed to neuron conversion (NC) medium based on DMEM:F12/Neurobasal (1:1) for three weeks. NC contains the following supplements: N2 supplement, B27 supplement (both 1x; Thermo Fisher Scientific), doxycycline (2 µg/ml, Sigma Aldrich), Laminin (1 µg/ml, Thermo Fisher Scientific Scientific), dibutyryl cyclic-AMP (100 µg/ml, Sigma Aldrich), DMH-1 (5 µM; Tocris), LDN-193189 (500 nM; Fisher Scientific Co) and A83-1 (500 nM; Santa Cruz Biotechnology Inc.), CHIR99021 (3 µM, LC Laboratories), Forskolin (5 µM, LC Laboratories) SB-431542 (10 µM; Cayman Chemicals), pyrintegrin (1 µM; Tocris), ZM336372 (.175 µM; Cayman), AZ960 (0.1 µM; Cayman), and KC7F2 (7.5 µM; Fischer Scientific). Medium was changed every third day.

### Flow Cytometry

For isolation of iNs and iPSC-iNs from 3 week cultures, cells were detached with TrypLE (Thermo Fisher) and stained for PSA-NCAM directly conjugated to APC (Miltenyl Biotec, 1x) for 1 hour at 4 C in sorting buffer (250mM myo-inositol and 5 mg/ml polyvinyl alcohol in PBS) containing 1% KOSR (Thermo Fisher). Cells were washed and resuspended in sorting buffer containing EDTA and DNAse and filtered using a 40-um cell strainer. For p16 flow analysis, cells were first fixed and permeabilized with 4% PFA and 0.5% Triton X-100. Primary antibodies (p16INK4a Abcam 1:250, TuJ Covance 1:3000) were applied overnight at 4C. Following two 10 minute washes with TBS, secondary antibodies (Donkey Anti-Mouse IgG Alexa Fluor 488/555 and Donkey Anti-Rabbit IgG Alexa Fluor 488/555, both 1:500) were incubated for 2 hours at room temperature. Cells were then washed twice and prepared for flow cytometry as described above.

### Whole Genome mRNA-seq and Analysis

Total bulk RNA was extracted from fibroblasts and iPSCs and from flow cytometry-isolated iNs and iPSC-iNs following 3 weeks of conversion using Trizol LS reagent (Thermo Fischer), followed by TURBO DNase digestion (Agilent). RNA integrity was assessed before library preparation using the TruSeq Stranded mRNA Sample Prep Kit according to the manufacturer’s instructions (Illumina). Libraries were sequenced paired-end 125 bp using the Illumina HiSeq 2500 platform. Read trimming was performed using TrimGalore, read mapping was performed using STAR, raw counts were generated using HOMER, variance stabilizing transformation normalization (vst) and differential expression analyses were performed in DESeq2. Statistical values were corrected for false discovery rates (FDR) using the Benjamini-Hochberg method implemented in R. Transcripts per million for all samples were generated by RSEM using standard paired-end or single-end settings as appropriate.

#### Genome-Wide DNA Methylation Analysis

Genomic DNA was extracted from flow cytometry-isolated iNs or bulk fibroblast cultures using the DNEasy Blood and Tissue Kit (Qiagen). DNA methylation assays were performed on the Illumina MethylEPIC BeadChip as per the standard manufacturers protocol. Raw IDAT files were processed and analyzed in R using the ChAMP and RnBeads packages, and normalized using the BMIQ procedure. Beta values were used as methylation residues for downstream analysis and correlation with RNAseq datasets.

Whole Genome assay for transposase-accessible chromatin using sequencing (ATACseq) and analysis ATAC-Seq was performed as described earlier (Buenrosto 2013). Briefly, 50,000 iNs were lysed in 50 ul lysis buffer (10 mM Tris-HCl ph 7.5, 10 mM NaCl, 3 mM MgCl2, 0.1% IGEPAL, CA-630, in water), pelleted and resuspended in 50 μL transposase reaction mix (1x Tagment DNA buffer, 2.5 μL Tagment DNA enzyme I in water (Illumina)), and incubated at 37°C for 30 min. DNA was purified with Zymo ChIP DNA concentrator columns (Zymo Research). DNA was then amplified with PCR mix (1.25 μM Nextera primer 1, 1.25 μM Nextera primer 2-bar code, 0.6x SYBR Green I (Life Technologies, S7563), 1x NEBNext High-Fidelity 2x PCR MasterMix, (NEBM0541) for 7-10 cycles, run on an agarose gel for size selection of fragments (160-500 bp), and extracted from the gel and paired-end 75 bp sequencing using the Illumina NextSeq 500 platform. Reads were trimmed using TrimGalore and mapped to the UCSC genome build hg38 using STAR. Peaks were then calculated and annotated using the HOMER software package. For agnostic genome-wide characterization of the peaks, sequence depth-normalized bigWig files were used for generating chromatin accessibility profiles using deepTools software. Next, differential accessibility analysis of all identified ATAC-Seq peaks was performed using differential peaks function in HOMER, and fold changes and significance of the resulting differential peaks were plotted as volcano plot using R ggplot2. Genome-wide integration of the ATAC-Seq data with the RNA-Seq data on a gene-by-gene level was performed based on comparing the fold changes of differentially accessible peaks (HOMER differential ATAC peaks) with the fold changes of differentially expressed genes (RNA-Seq expression), using R, and visualization was performed using R ggplot2 and GraphPad Prism software. GSEA of annotated Promoter-TSS peaks was performed using STRING, and overlapping with the Riessland et al dataset was performed using shared GO terms and enrichment scores present in both datasets. Motif enrichment analysis was performed in HOMER. Statistical enrichment of motifs was performed by using all control iNs as background for AD iN peak and subsequent motif calling. The resulting q-values generated through this method are marked in the figures. Comparisons of peak height between AD and CTL iNs was performed in deeptools using the plotHeatmap function.

### SunTag Labeling of L1Hs DNA

A lentiviral SunTag reporter construct was obtained from VectorBuilder which encoded nuclease-dead Cas9 (dCas9) fused to tandem GCN4 peptide repeats and a nuclear export signal (NES; MTKKFGTLTI). Cells were transduced with the SunTag reporter lentivirus and selected for using G418 (Gibco). A separate lentiviral vector was used to deliver a gRNA targeting the 5′ untranslated region of human-specific LINE-1 elements (L1Hs, TGGTGCGCCGTTTCTTAAGC) with scFv-sfGFP fragments targeting GCN4. Control cells received a non-targeting gRNA scramble. High content imaging was performed using a Yokogawa imaging system on either live cells or PFA fixed and stained cells as indicated. Nuclear regions were segmented with DAPI for quantification of cytoplasmic fluorescence.

### EdU Labeling

For EdU labeling, induced neurons were cultured on ibidi imaging plates (80826) as previously described and treated with the indicated LINE-1 modulating compounds for 48 hours (3TC, FTC) or 1 week (ASO). Following treatment, EdU was added directly to the culture medium at a final concentration of 10 uM for eight hours to label newly synthesized DNA. Following EdU incorporation, cells were washed with PBS and fixed with 3.7% formaldehyde in PBS for 15 minutes, followed by 0.5% Triton X-100 permeabilization for 20 minutes. EdU incorporation was fluorescently labeled by click-it chemistry (Invitrogen C10337), and cells were stained for mitochondria (TOM20), a neuronal marker (MAP2), and DAPI as described below. A TOM20 based mask was applied in fiji to exclude mitochondrial EdU signal, and EdU positive cytoplasmic puncta were quantified per patient from three separate fields.

### Transposable Element and LINE-1 Expression Quantification

To quantify transposable element (TE) and LINE-1 expression at single-cell resolution, single-cell PacBio Kinnex libraries were generated from samples obtained from 9 individuals, including 4 Alzheimer’s disease cases, 3 controls, and 2 individuals with Alzheimer’s disease-related dementia using the manufacturers specifications. Libraries were sequenced using PacBio long-read sequencing, yielding 5,636,423 HiFi reads and 83,674,058 full-length cDNA reads. Filtered reads were aligned to the human reference genome (GRCh38, Ensembl release 115) using STARlong (v2.7.11b) with parameters permitting multimapping reads and retaining single-cell barcode/UMI-related tags (--outFilterMultimapNmax 100, --winAnchorMultimapNmax 100, --seedPerReadNmax 10000). Cell barcodes were matched against the 10x Genomics 3M 3′ gene expression May 2023 reverse-complement whitelist. TE and LINE-1 expression was quantified using iRescue (v1.1.2) with the merged annotated BAM. Three separate quantification runs were performed per patient using custom region BED files: (1) all RepeatMasker-annotated TE families (GRCh38, downloaded from UCSC); (2) 138 active full-length L1HS loci from L1Base2, filtered to remove elements lacking ORF2 or classified as inactive; and (3) 13,292 inactive L1 loci from L1Base2, filtered to remove elements with ORF2 or classified as active. Count matrices were loaded into Seurat (v5.3.0) alongside protein-coding gene expression from the standard DGE output.

### Spatial Transcriptomics Analysis

Publicly available post-mortem human brain spatial transcriptomics datasets from the ST011 series of ssRead archive (bmblx.bmi.osumc.edu/ssread, GEO GSE220442) were analyzed, including three control samples and three mid-stage Alzheimer’s disease samples. L1HS quantification was performed using the SoloTE Seurat analysis pipeline using default parameters for 10X single cell input. Control and AD samples were merged separately using shifted spatial coordinates, and Delaunay spatial networks were generated for each merged diagnostic group. Spatial enrichment of p16-positive and L1HS-high cells was assessed by calculating the fraction of Delaunay edges connecting CDKN2A/p16-positive cells to Active L1HS-high cells. Statistical significance was estimated using 1,000 permutations in which p16 labels were randomly shuffled across cells, generating empirical null distributions, z scores, and permutation-based p values.

For each cell/spot, an inflammatory module score was calculated as the mean normalized expression of inflammatory and interferon-response genes, IL1B, TNF, IL6, CXCL10, CCL2, CCL3, CCL4, STAT1, IRF1, ISG15, IFIT1, MX1, CD74, and HLA-DRA. To quantify local inflammatory context, each cell was then assigned a neighbor inflammatory score equal to the average inflammatory module score of its directly connected spatial neighbors in the Delaunay network.

### Astrocyte Culture

Human cortical astrocytes were acquired from a commercial vendor and cultured in astrocyte medium according to the manufacturer’s specifications (ScienCell 1800, 1801). For CM experiments, 48-hour iN conditioned supernatant from three 3TC treated AD lines was spiked in at a 1:2 ratio with untreated astrocyte medium for 48 hours. Protein input between samples was normalized to total protein as measured by qubit (ThermoFisher). Cells were then fixed for ICC or harvested with Trizol for RNA extraction.

### Immunofluorescence

Cells were cultured on tissue culture-treated ibidi μ-slides for imaging. Cells were fixed with 4% PFA for 20 min at room temperature and washed 3 × 15 minutes with TBS, followed by a 1-hr block with TBS containing 10% serum and 0.1% Triton X-100. Primary antibodies (p16INK4a Abcam 1:250, TuJ Covance 1:3000, NeuN EMD Millipore 1:250, GFAP EMD Millipore 1:1000, TOM20 Millipore 1:500, GM130 Cell Signaling 1:3000, STING Thermo Fischer 1:500, MAP2 abcam 1:5000) were applied overnight at 4C. Following two 10 minute washes with TBS, nuclei were stained with DAPI (1:10,000 Sigma-Aldrich) and secondary antibodies (Donkey Anti-Mouse IgG Alexa Fluor 488/555 and Donkey Anti-Rabbit IgG Alexa Fluor 488/555, and Donkey Anti-Chicken IgG Alexa Fluor 488/555, all 1:250) were incubated for 2 hours at room temperature. After washing, slides were mounted in PVA-DAPCO (Sigma Aldrich). Confocal images were taken on standard fluorescence microscopes, Zeiss LSM880 confocal, or Yokogawa spinning disk microscopes.

ImageJ software analyze particles was used for regions of interest (ROI) selection and quantification of immunofluorescent signals within ROIs. Proximity analysis was performed with 3D ImageJ Suite. All data for one experiment were acquired from cells cultured or tissue processed in parallel on the same microscope with the exact same setting used.

